## Supplementary Fig for "Genomic signatures of somatic hybrid vigor due to heterokaryosis in the oomycete pathogen, *Bremia lactucae*"

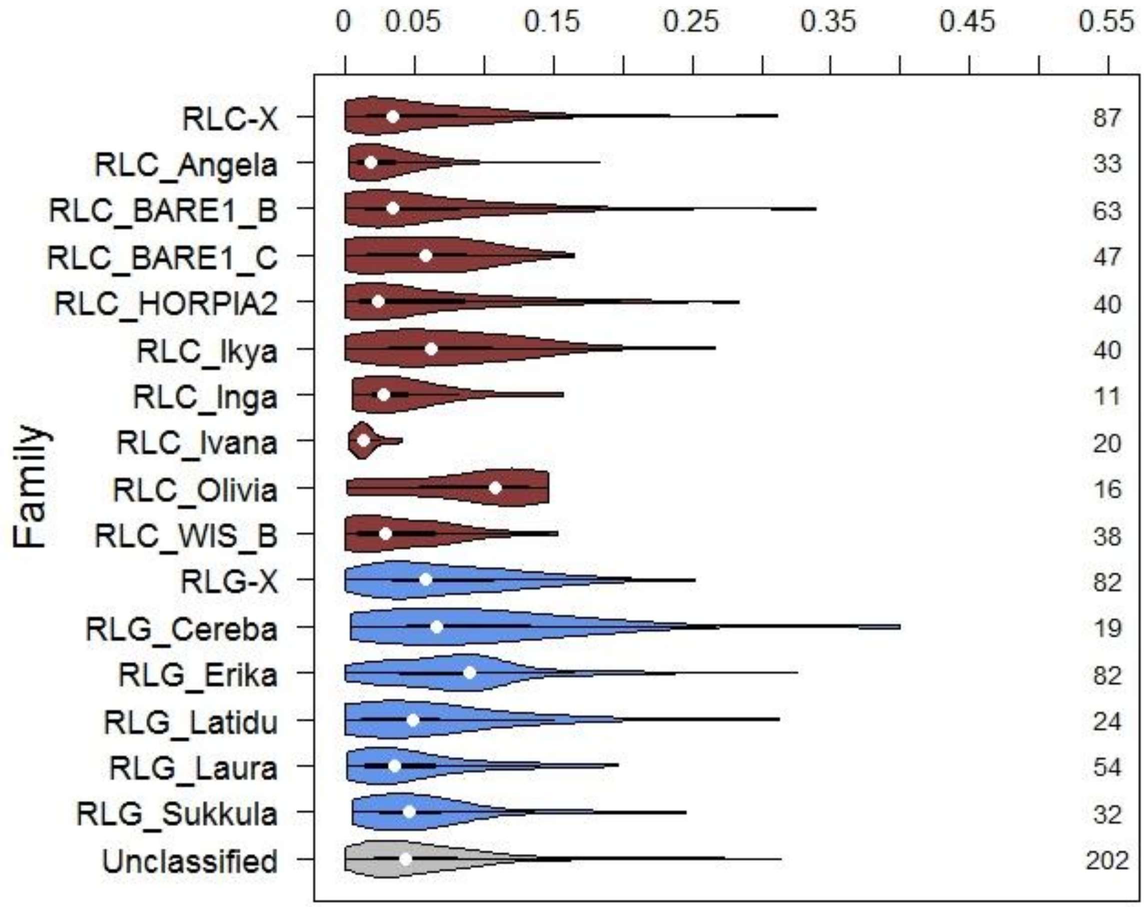


**Supplementary Figure 1. Long tandem repeat (LTR) nucleotide divergence.**

Estimates of divergence between LTRs identified in the *B. lactucae* assembly. LTRs are identical upon insertion of an LTR retrotransposon. Their divergence can therefore be used to infer the timing of insertion.  RLC (red) = LTR *Copia* elements. RLG (blue) = LTR *Gypsy* elements.

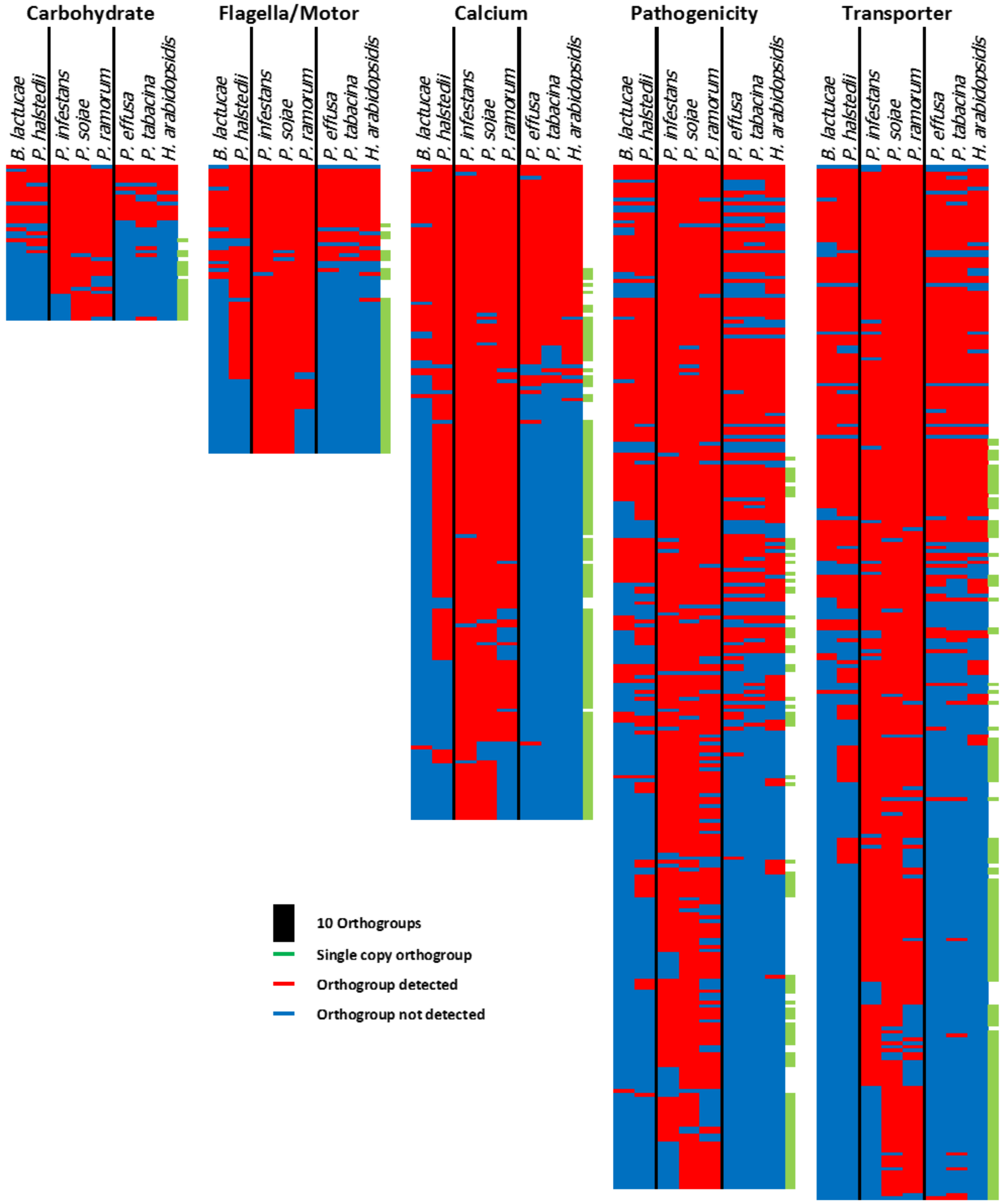


**Supplementary Figure 2. Heat map showing the distribution of orthogroups where a Pfam domain, found to be depleted in downy mildews, is detected as encoded by at least one gene model of that orthogroup.**

Red indicates the orthogroup contains at least one model from the species, blue indicates that no model is detected from the species. Green tabs on the right of the row indicates that the orthogroup contains a maximum of one gene from each species except for *P. effusa* and *P. tabacina* where two models were permitted because two isolates were combined in the analysis.

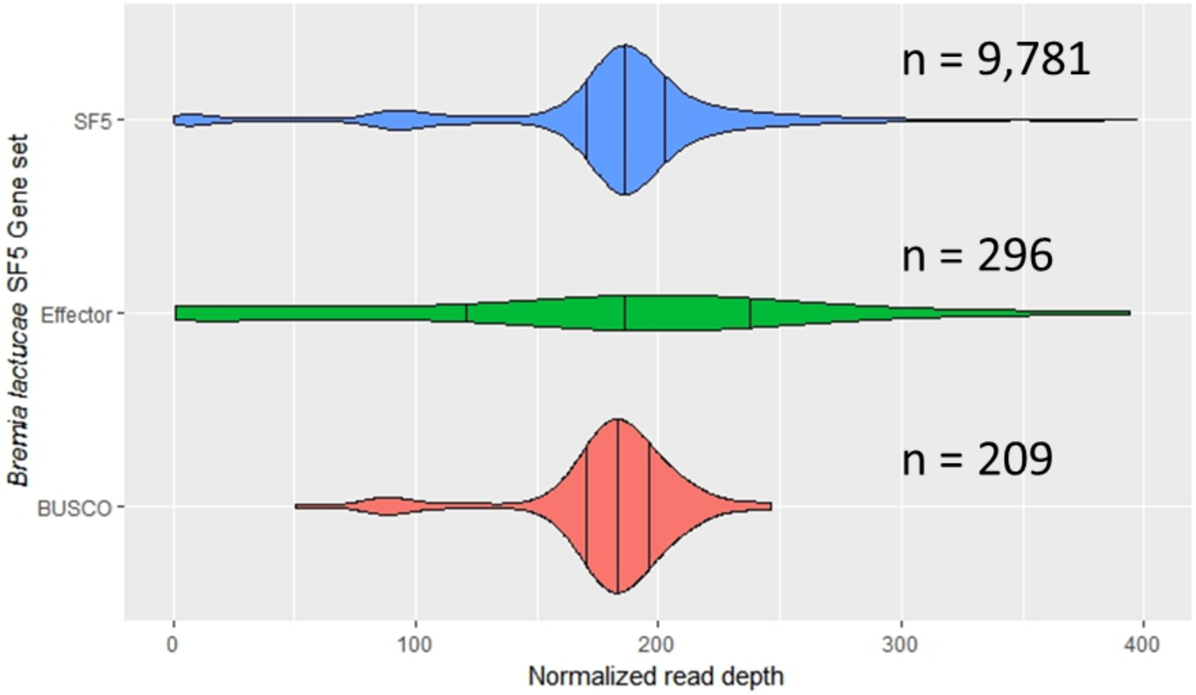


**Supplementary Figure 3. Sequence coverage of different groups of genes in the *B. lactucae* assembly.**

The majority of annotated genes have coverage at approximately 180x, equal to the sequencing depth. This indicates that the majority of genes are assembled as single consensus sequences. A minority of genes have coverage at approximately 90x. This is consistent with alleles being assembled independently or hemizygous regions. The same pattern is observed in the BUSCO genes, of which eight were detected as being duplicated. The effector portion of the genomes does not display the same distribution. This may be due to effector alleles being assembled independently, or alternatively a high rate of divergence between haplotypes resulting in poor rates of mapping.

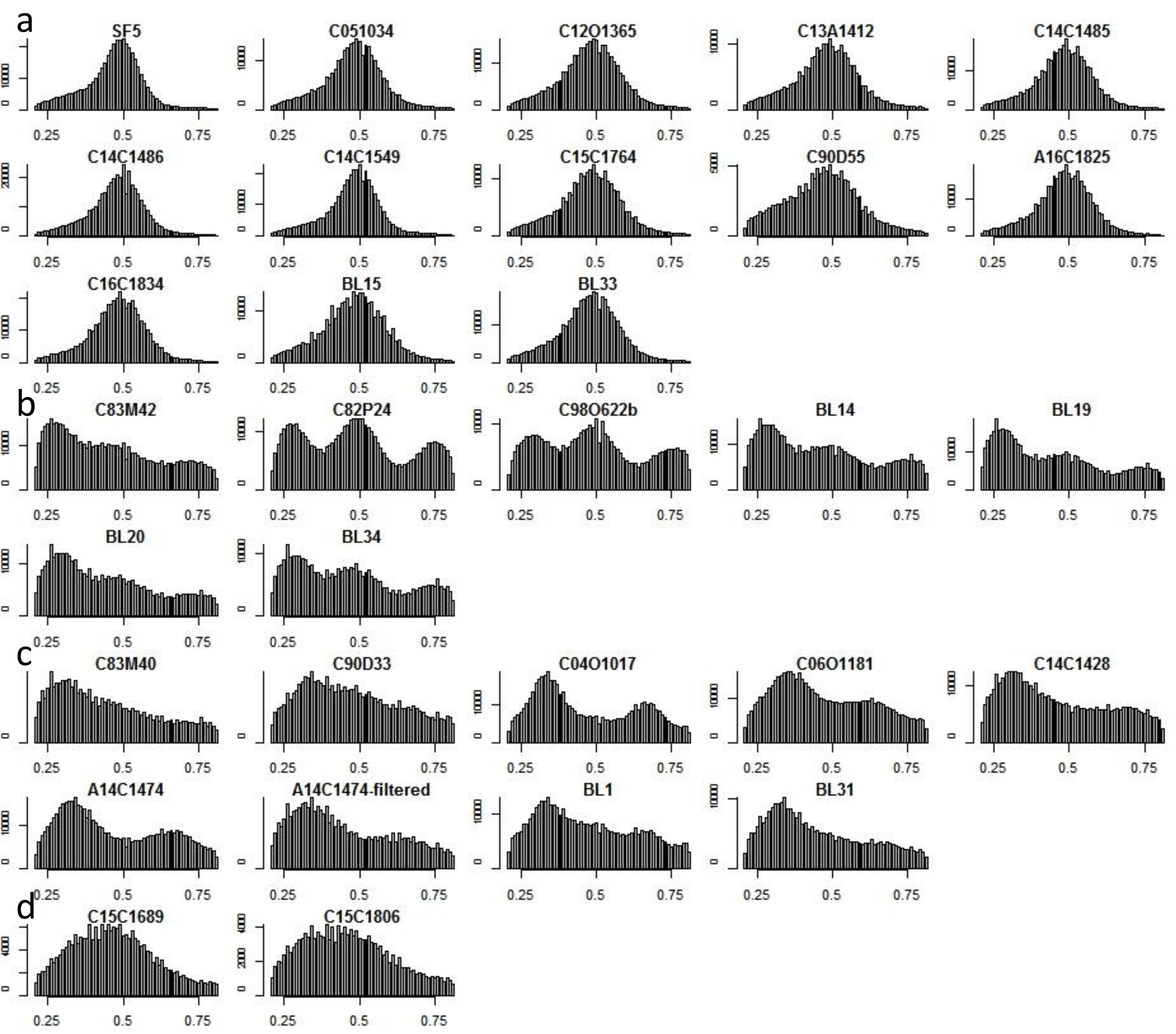


**Supplementary Figure 4. Bar plots of alternative allele frequencies for each SNP detected between the tested isolate and the SF5 assembly.**

The x axis defines the frequency of reads for each SNP. The y axis defines the number of SNPs with that frequency. a) Ten American isolates and three European isolates, including the reference isolate SF5 with a unimodal distribution with a peak at 0.5 indicative of a homokaryotic state. b) Three Californian and four European isolates with a trimodal distribution with peaks at 0.25, 0.5, and 0.75 consistent with a heterokaryotic with two nuclei in a 1:1 ratio. c) Six American isolates, including one isolate which was filtered (A14C1474-filtered) and two European isolates with bimodal distributions with peaks at 0.33 and 0.67 consistent with a heterokaryon made up of three nuclei in a 1:1:1 ratio. d) Two Californian isolates had SNP frequency profiles which did not have clear unimodal, trimodal, or bimodal distributions; these could be heterokaryons containing multiple nuclei.

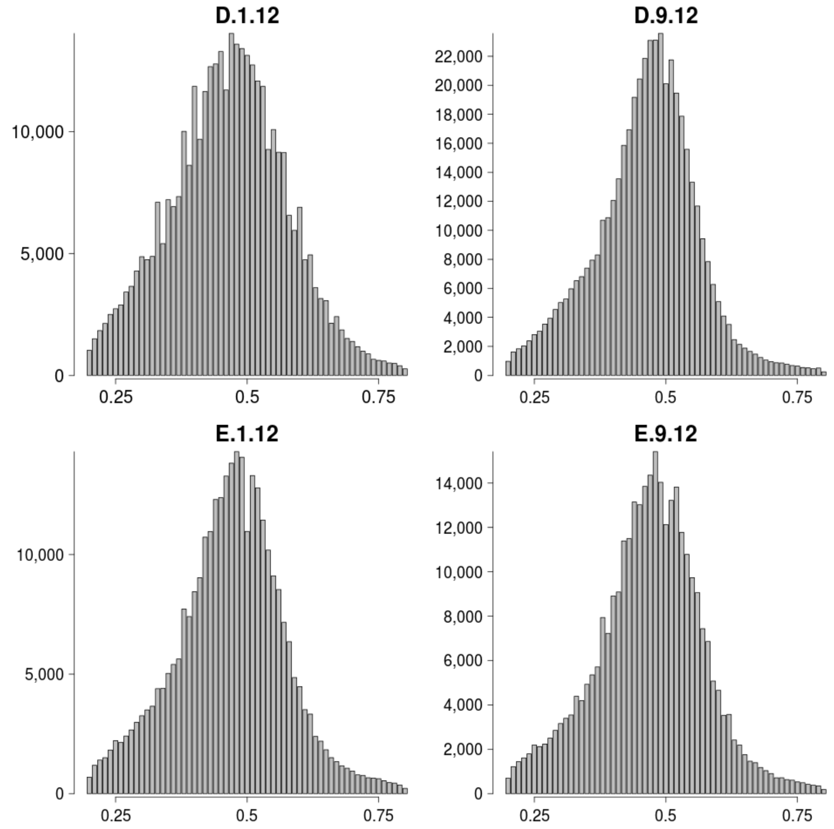


**Supplementary Figure 5. Bar plots of alternative allele frequencies for each SNP detected between 4 progeny isolates and the SF5 assembly.**

All 4 sexual progeny from the SF5 x C82P24 cross tested had a unimodal distribution with a peak at 0.5. This is consistent with each sexual progeny being a diploid, with a single haploid gamete being contributed from each parent.

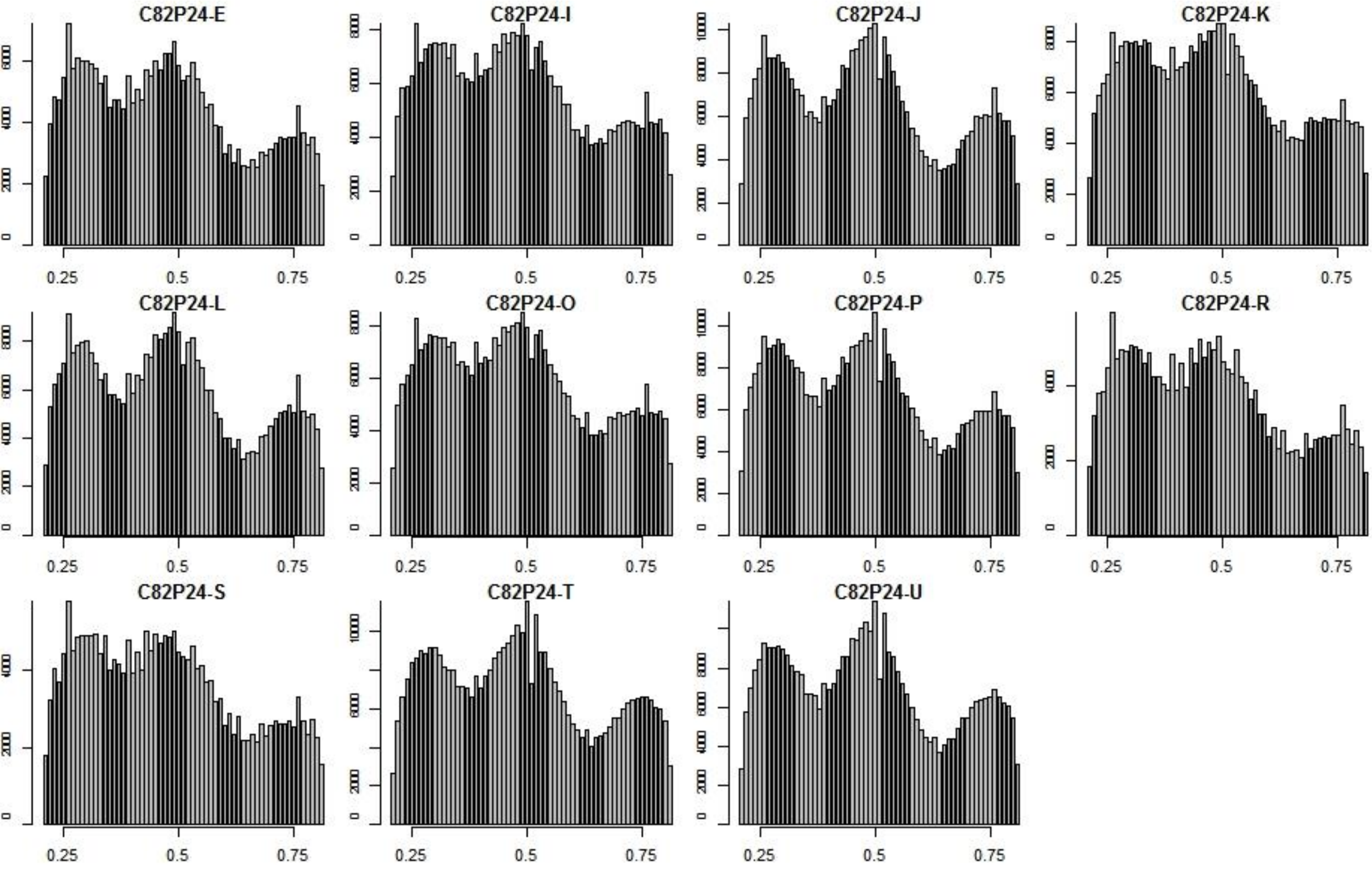


**Supplementary Figure 6. Bar plots of alternative allele frequencies for each SNP detected between 11 C82P24 single spore derivatives and the SF5 assembly.**

All 11 derivatives had a trimodal distribution with three peaks at 0.25, 0.5 and 0.75 consistent with each multinucleate spore transmitting two genetically distinct nuclei to the next asexual generation.

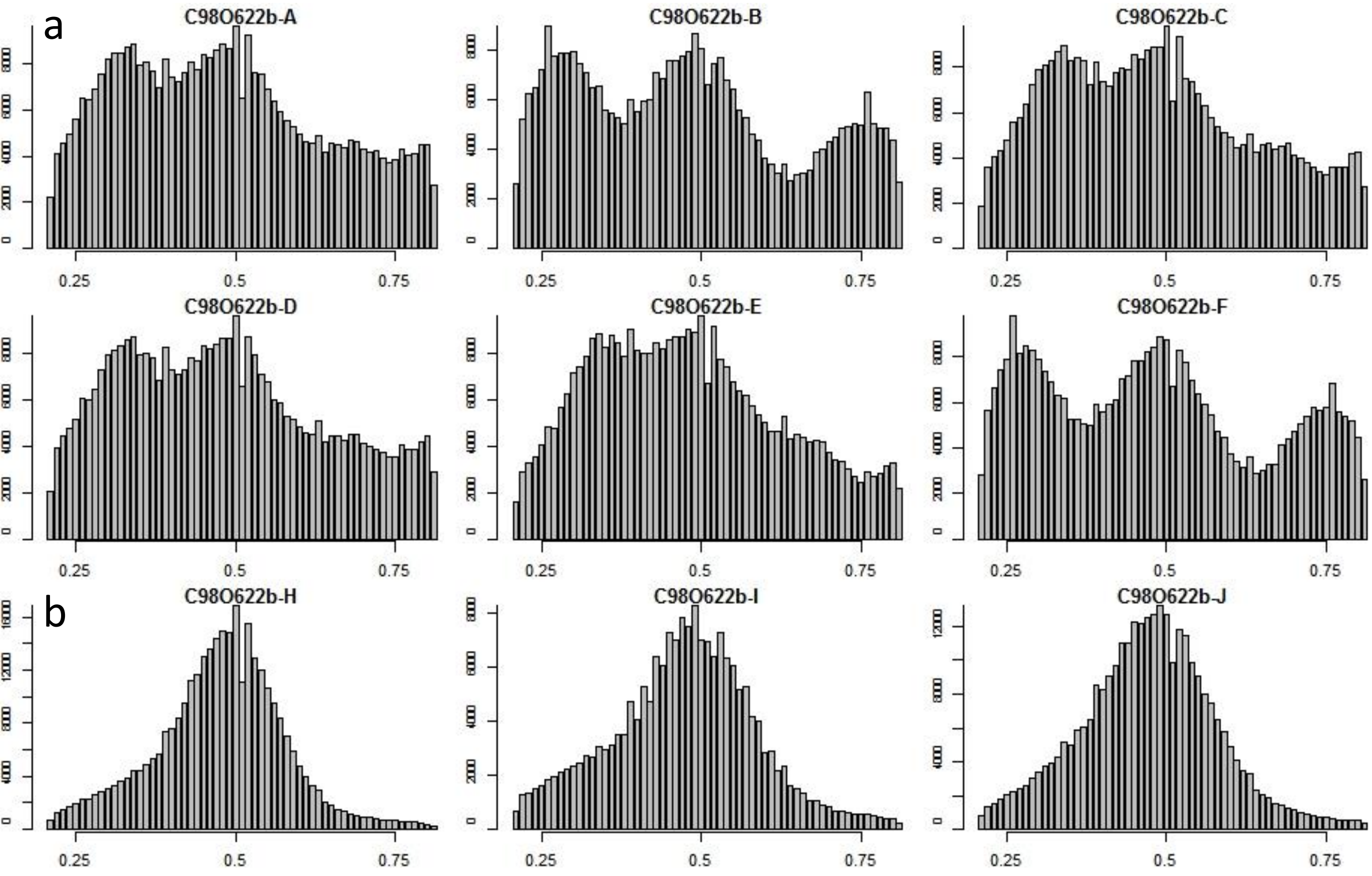


**Supplementary Figure 7. Bar plots of alternative allele frequencies for each SNP detected between nine C98O622b single spore derivatives and the SF5 assembly.**

a) Six of the nine derivatives had trimodal distributions; two (B and F) had peaks at 0.25, 0.5, and 0.75, while four (A, C, D & E) had peaks at 0.33, 0.5, and 0.66. b) Three of the nine derivatives had a unimodal distribution with a single peak at 0.5 consistent with the transmission of a single nuclear type through their progenitor single spores.

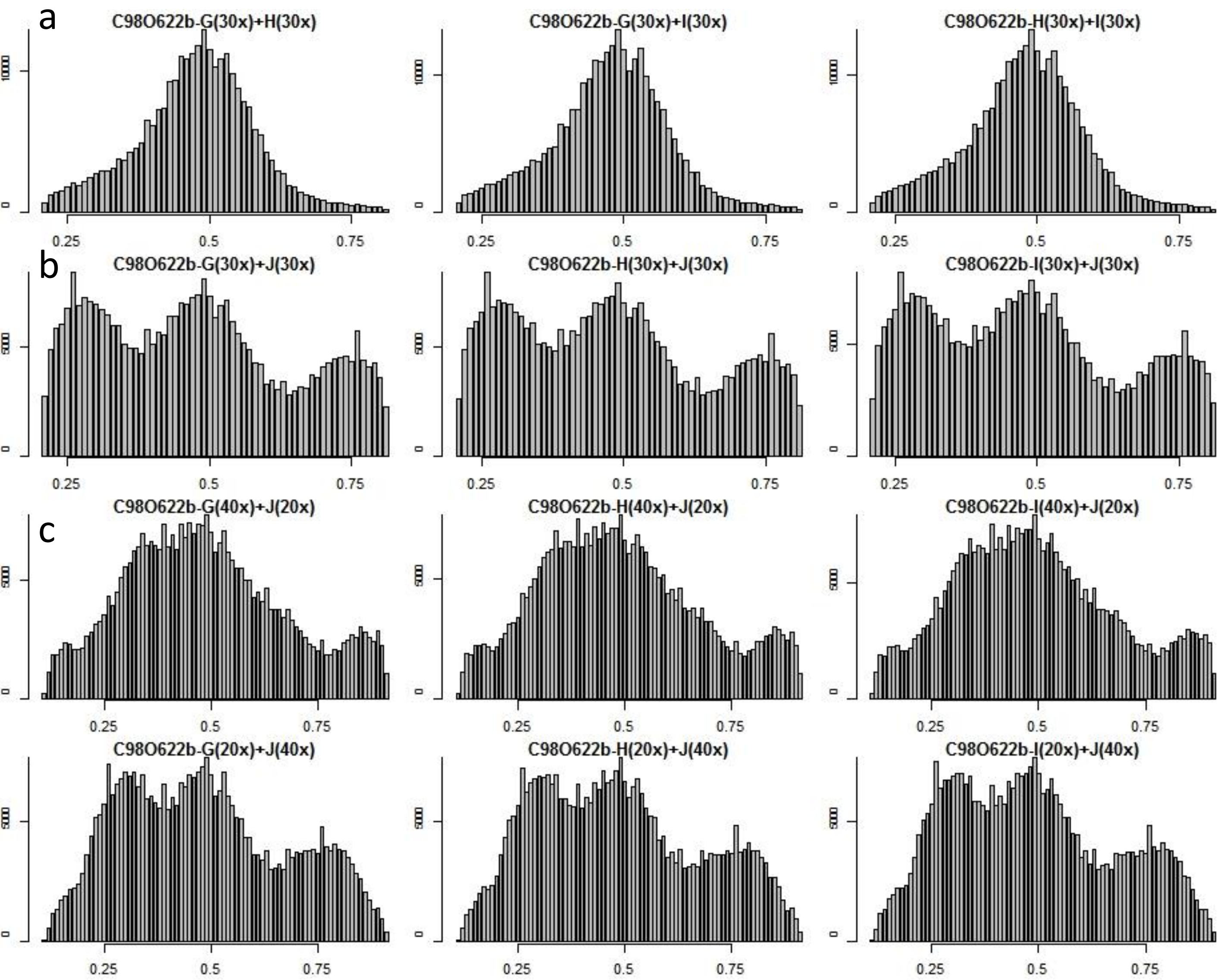


**Supplementary Figure 8. Bar plots of alternative allele frequencies for each SNP detected between heterokaryons generated *in silico* and the SF5 assembly.**

a) Combining reads from two homokaryons with high identity to one another produced a single peak, consistent with them being highly similar isolates, as demonstrated by kinship analysis (Figure 6). b) Combining reads from two different homokaryon nuclear types in equal combinations, simulating a 1:1 ratio produced plots with three peaks at approximately 0.25, 0.5 and 0.75, similar to C98O622b heterokaryons B and F (Figure 6, Supplementary Figure 6). c) Combining reads in 2:1 or 1:2 proportions between homokaryon nuclear types resulted in plots similar to C98O622b heterokaryons A, C, D & E (Figure 6, Supplementary Figure 6).

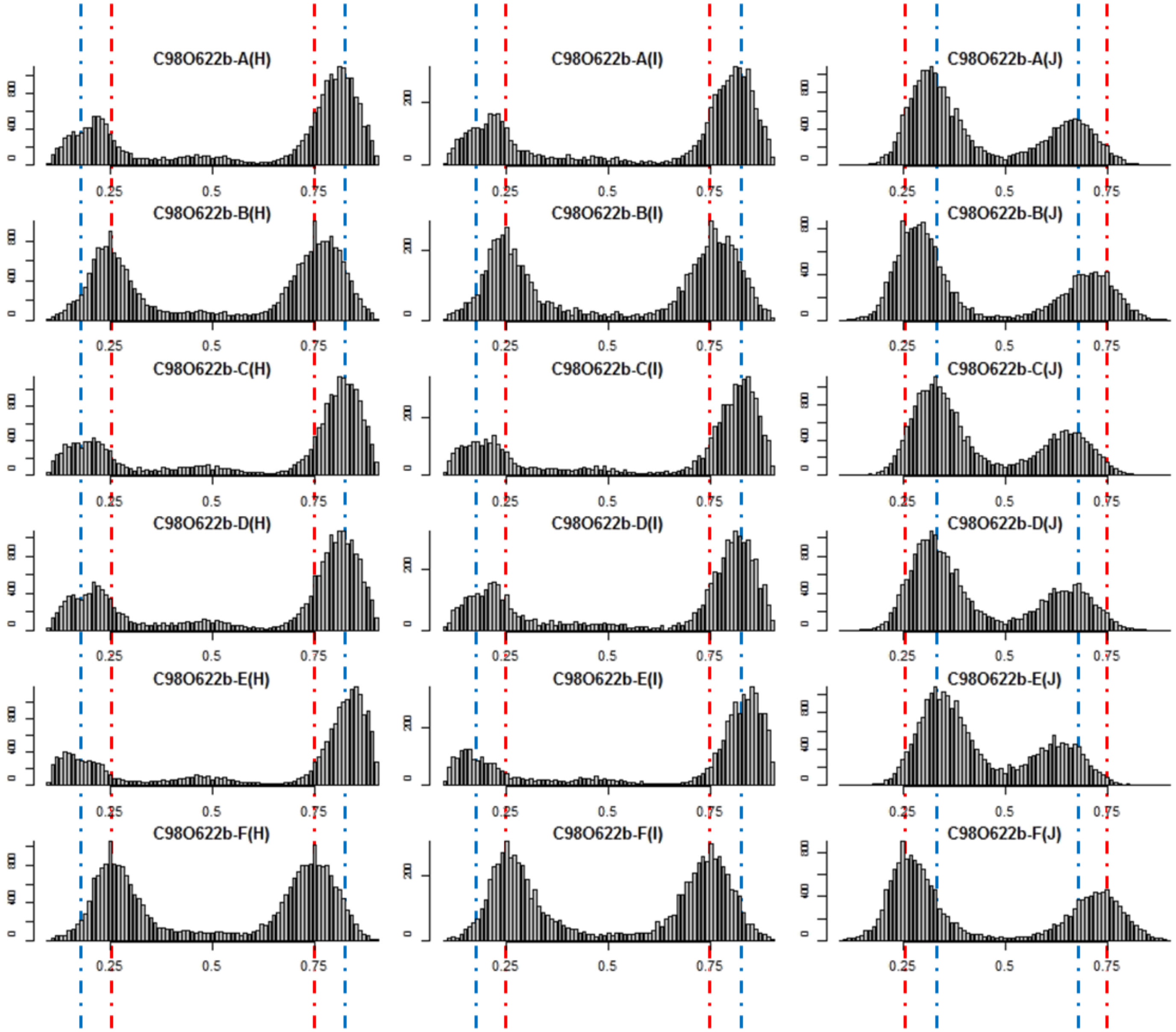


**Supplementary Figure 9. Bar plots of alternative allele frequencies for SNPs unique to each homokaryotic nuclear type in the heterokaryotic derivatives and the SF5 assembly.**

SNPs unique to either homokaryotic derivative nuclear type (H/I or J) were sampled from all six heterokaryotic derivatives of C98O622b. In heterokaryotic derivatives B and F (second and sixth row) with peaks previously at 0.25, 0.5 and 0.75 (Figure 6c, Supplementary Figure 7), all SNPs were detected at approximately 0.25 and 0.75 (indicated by red dotted lines). This is consistent with genetically distinct nuclei being present in a 1:1 ratio. In heterokaryons A, C, D & E, where peaks were not at 0.25, 0.5 and 0.75 (Figure 6c, Supplementary Figure 7), SNPs unique to derivative H or I (column two) were consistently at 0.17 and 0.83 (blue dotted line) while SNPs unique to J (column three) were at 0.33 and 0.66 (blue dotted line). This pattern is consistent with the second nucleus being present twice as often as the first nucleus in these heterokaryotic derivatives.

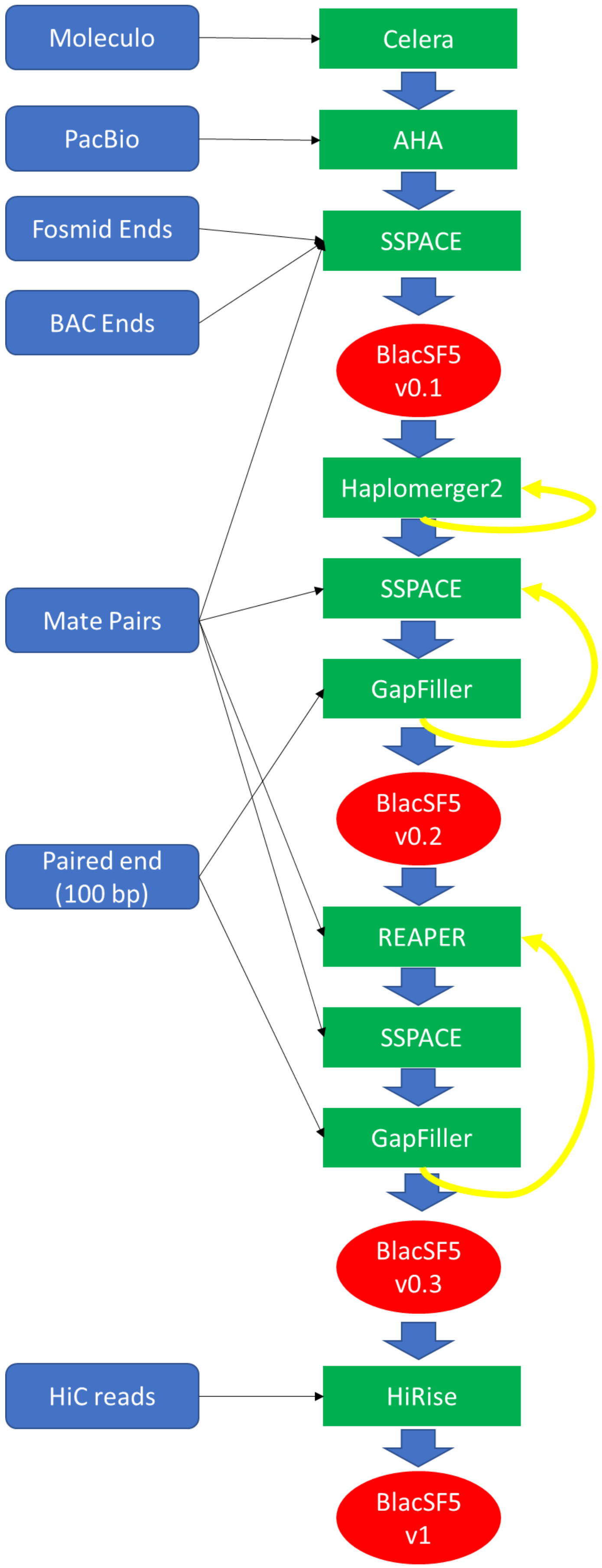


**Supplementary Figure 10a. Assembly workflow used to achieve the final *B. lactucae* SF5 assembly.**

Blue boxes are input data. Green boxes are software used to incorporate the data. Red circles are checkpoints assessed in Supplementary Figure 10b. Black arrows connect input data to the software used. Blue arrows show the linear progression through the workflow. Yellow arrows indicate where software(s) was applied iteratively until minimal changes were detected between input and output assembly statistics.

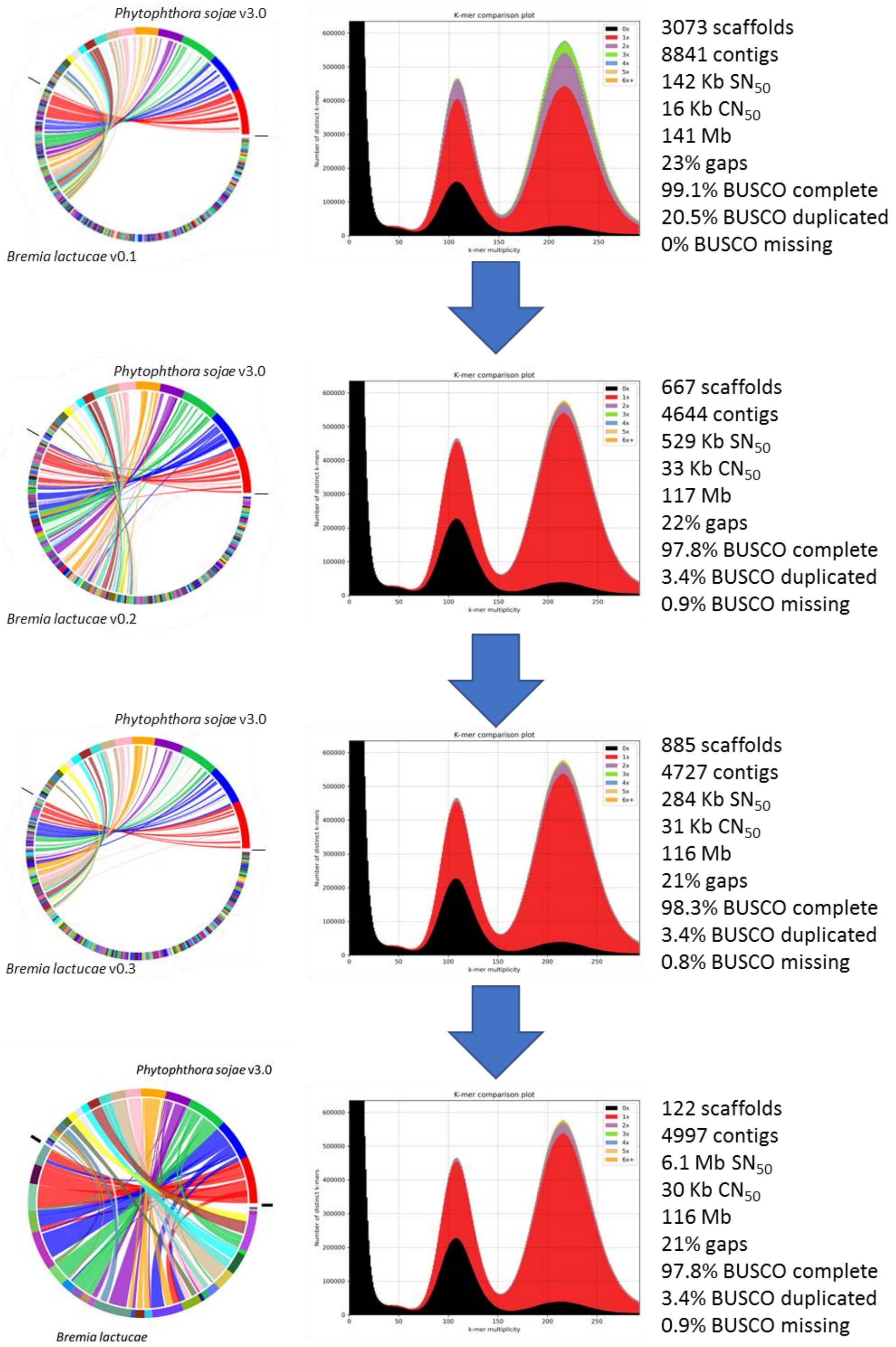


**Supplementary Figure 10b. Genome assembly checkpoints.**

SyMap plots, displaying nuclear collinearity with *P. sojae* v3.0 became more dense as the scaffold $N_{50}$ increased. HiRise significant improved collinearity. KAT spectra-cn plots were used to infer redundancy within the assembly. The initial assembly had high levels of duplication, which were removed by Haplomerger2. Assembly statistics at each checkpoint are shown to the right.
