## Supplementary Table for "Genomic signatures of somatic hybrid vigor due to heterokaryosis in the oomycete pathogen, *Bremia lactucae*"

**Supplementary Table 1. Total nuclear DNA content of 39 *Bremia lactucae* isolates, including SF5. Sexual progeny isolates generated in the lab are italicized. Table is sorted by mean nuclear content (Mb).**

| Isolate label | Mean nuclear content | Standard Deviation | Replicates |
|---|---|---|---|
| *A.2.12* | *312.5* | *7.8* | *2* |
| *A14C1474* | *311.5* | | *1* |
| *CrAIII-19* | *311.5* | *5.0* | *2* |
| C15C1816 | 310.5 | | 1 |
| *Se6-2* | *310.1* | *0.9* | *2* |
| C08O1181 | 309.6 | 3.7 | 2 |
| SF5 | 307.9 | 4.2 | 16 |
| A14C1435 | 307.8 | 3.8 | 4 |
| *CrAIII-18* | *307.8* | *4.8* | *2* |
| *A.5.12* | *307.4* | *0.7* | *2* |
| *V1* | *306.8* | *6.6* | *2* |
| C04O1017 | 306.6 | 2.9 | 2 |
| C05R1034 | 306.0 | 1.6 | 3 |
| *B.9.12* | *305.8* | *2.5* | *2* |
| A16C1821 | 305.6 | | 1 |
| C82P24 | 305.4 | 2.4 | 18 |
| *ZL07A* | *305.3* | | *1* |
| C15C1806 | 305.2 | | 1 |
| *C.7.12* | *305.0* | | *1* |
| *CrAII-2* | *304.8* | | *1* |
| C98O648 | 304.5 | 16.0 | 3 |
| C98O622 | 304.4 | 7.0 | 3 |
| *A.17.12* | *304.2* | *2.7* | *2* |
| *CrA III-2* | *303.6* | *2.0* | *2* |
| C15C1771 | 303.3 | 3.8 | 17 |
| C13C1414 | 302.8 | 8.5 | 3 |
| C91D39 | 301.8 | 1.9 | 2 |
| *CrAIII-15* | *301.6* | *6.9* | *2* |
| C83M47 | 301.3 | 1.7 | 2 |
| C14C1549 | 301.0 | 5.2 | 2 |
| C14C1485 | 300.7 | 3.7 | 4 |
| C16C1832 | 300.6 | | 1 |
| C14C1452 | 298.9 | 9.1 | 3 |
| C15C1769 | 298.9 | 4.2 | 6 |
| C98O622b | 296.9 | 8.6 | 3 |
| C13A1407 | 294.9 | 7.4 | 3 |
| C14C1486 | 294.5 | 3.4 | 6 |
| C15C1622 | 291.3 | 12.6 | 4 |
| *R7* | *288.8* | *8.1* | *5* |
| **Mean** | 303.8 | 5.4 | |

**Supplementary Table 3. Orthology of previously reported candidate effectors of *B. lactucae* to proteins annotated in the SF5 genome.**

| Orthogroup | Stassen *et al.* 2013 / Giesbers *et al.* 2017 / Pelgrom *et al.* 2018 | SF5 assembly (this paper) |
|---|---|---|
| 1 | BLR17 | BlacSF5_002585-RA |
| | BLR28 | BlacSF5_003650-RA |
| | BLR33 | BlacSF5_009425-RA |
| 2 | BLR19 | BlacSF5_006115-RA |
| | **BLR31** | BlacSF5_008983-RA |
| | **BLR40** | |
| 3 | BLQ01 | BlacSF5_008167-RA |
| | | BlacSF5_008168-RA |
| | | BlacSF5_008173-RA |
| | | BlacSF5_009528-RA |
| 4 | BLG02 | BlacSF5_002342-RA |
| | **BLG03** | BlacSF5_004206-RA |
| 5 | BLR09 | BlacSF5_007121-RA |
| | BLR10 | BlacSF5_007131-RA |
| 6 | BLR11 | BlacSF5_004761-RA |
| | BLR35 (3) | BlacSF5_004760-RA (3) |
| 7 | BLR20 (1) | BlacSF5_003745-RA |
| | | BlacSF5_003751-RA |
| | | BlacSF5_003753-RA |
| 8 | **BLN08 (4)** | BlacSF5_000769-RA (2) |
| | BLR36 (5) | BlacSF5_003408-RA (3) |
| 9 | BLN01 (7) | BlacSF5_008217-RA (7) |
| | **BLN06 (6)** | BlacSF5_008226-RA (5) |
| 10 | BLR05 | BlacSF5_006000-RA |
| | | BlacSF5_006001-RA |
| 11 | BLR15 | BlacSF5_005890-RA |
| | BLR30 | |
| 12 | BLR16 | BlacSF5_005893-RA |
| | | BlacSF5_009559-RA |
| 13 | BLR18 | BlacSF5_002279-RA |
| | | BlacSF5_002280-RA |
| 14 | BLR25 | BlacSF5_004523-RA |
| | | BlacSF5_006920-RA |
| 15 | BLR32 | BlacSF5_008977-RA |
| | | BlacSF5_008980-RA |
| 16 | **BLR38** | BlacSF5_005743-RA |
| | | BlacSF5_005744-RA |
| 17 | BLC01 | BlacSF5_001066-RA |
| | | BlacSF5_009260-RA |
| 18 | BLN05 | BlacSF5_002598-RA |
| | BLQ04 | |
| 19 | **BLG01** | BlacSF5_006965-RA |

| | | |
|---|---|---|
| 20 | BLR02 | BlacSF5_002587-RA |
| 21 | BLR06 | BlacSF5_002950-RA |
| 22 | BLR07 (4) | BlacSF5_008762-RA (3) |
| 23 | BLR13 (5) | BlacSF5_002525-RA (3) |
| 24 | BLR23 | BlacSF5_008952-RA |
| 25 | BLR27 | BlacSF5_002491-RA |
| 26 | BLR37 | BlacSF5_008452-RA (5) |
| No orthology in SF5 assembly | BLR01 | |
| | BLR03 | |
| | BLR04 | |
| | BLR08 | |
| | BLR12 | |
| | BLR14 | |
| | BLR21 | |
| | BLR22 | |
| | BLR24 | |
| | BLR26 | |
| | BLR29 | |
| | BLN03 | |
| | BLN04 | |
| | | 234 candidate effectors novel to SF5 assembly |

**Key:**

Red: Contains WY (# domains detected)

**Bold: Effector response detected**

| | |
|---|---|
| BLR | RxLR motif present |
| BLQ | QxLR motif present |
| BLN | RxLR not detected |
| BLC | Crinkler |

**Supplementary Table 4. Pathogenicity domains annotated in genome assemblies of Peronosporales spp.**

| | *Bremia lactucae* | *Peronospora* | | | | *Hyaloperonospora arabidopsidis* | *Plasmopara halstedii* | *Phytophthora* | | | Downy mildews *vs.* *Phytophthora* spp. | *Phytophthora* spp. + *P. halstedii* + *B. lactucae* *vs.* Peronosporales lineage |
|---|---|---|---|---|---|---|---|---|---|---|---|---|
| | | *effusa* | | *tabacina* | | | | | | | | |
| | SF5 | R13 | R14 | J2 | S26 | | | *sojae* | *infestans* | *ramorum* | **t-test** | **t-test** |
| Serine protease | 27 | 14 | 13 | 21 | 12 | 12 | 27 | 42 | 37 | 36 | | <0.05 |
| Aspartic protease | 11 | 11 | 9 | 17 | 7 | 15 | 26 | 147 | 20 | 116 | <0.05 | |
| Cysteine protease | 10 | 16 | 17 | 18 | 17 | 16 | 20 | 28 | 27 | 30 | | |
| Metalloprotease | 18 | 16 | 18 | 14 | 16 | 25 | 17 | 38 | 24 | 26 | | |
| Kazal-like serine protease inhibitor | 3 | 1 | 1 | 2 | 2 | 7 | 17 | 45 | 35 | 17 | <0.01 | <0.01 |
| Cystatin-like cysteine protease inhibitor | 1 | 1 | 1 | 1 | 1 | 1 | 3 | 4 | 6 | 4 | <0.01 | <0.05 |
| Cutinase | 0 | 0 | 0 | 0 | 0 | 2 | 2 | 15 | 4 | 4 | <0.01 | |
| Pectin lyase | 14 | 11 | 13 | 6 | 8 | 22 | 19 | 122 | 100 | 82 | <0.001 | <0.05 |
| CAP domain | 41 | 39 | 42 | 45 | 48 | 38 | 72 | 155 | 112 | 104 | <0.01 | |
| NPP1-like | 13 | 9 | 10 | 17 | 14 | 21 | 19 | 80 | 28 | 62 | <0.01 | |
| Elicitin-like | 20 | 16 | 15 | 12 | 8 | 20 | 20 | 77 | 54 | 61 | <0.001 | <0.05 |
| Jacalin | 15 | 10 | 10 | 7 | 7 | 10 | 20 | 31 | 22 | 27 | | <0.05 |
| Frequency | 0.0177 | 0.0168 | 0.0173 | 0.0141 | 0.013 | 0.0132 | 0.0167 | 0.0295 | 0.0262 | 0.0361 | <0.001 | <0.05 |

**Supplementary Table 5. Nuclear DNA content of 12 asexual single sporangia derivatives of *B. lactucae* isolate C82P24.**

| Derivative | Nuclear DNA content |
|---|---|
| P24 A | 304.2 |
| P24 B | 304.8 |
| P24 C | 302.0 |
| P24 D | 305.7 |
| P24 E | 308.5 |
| P24 F | 301.7 |
| P24 G | 301.3 |
| P24 H | 297.5 |
| P24 I | 301.8 |
| P24 J | 301.2 |
| P24 K | 308.9 |
| P24 L | 300.9 |
| **Mean** | 303.2 s.d. +/-3.3 |

**Supplementary Table 8. SRA accession numbers for oomycete sequences downloaded for this study.**

| **Genus** | **Species** | **Strain** | **SRA #** |
|---|---|---|---|
| *Albugo* | *candida* | Nc2 | SRR1811450, SRR1811464 |
| *Albugo* | *candida* | BoL | SRR1811474 |
| *Albugo* | *candida* | BoT | SRR1811472, SRR1811473 |
| *Albugo* | *candida* | 2v | SRR1811471 |
| *Albugo* | *candida* | EM2 | SRR1806791 |
| *Albugo* | *laibachii* | Nc14 | ERR022981, ERR022982 |
| *Aphanomyces* | *astaci* | APO3 | SRR957817 |
| *Aphanomyces* | *invadans* | NJM9701 | SRR957818 |
| *Hyaloperonospora* | *arabidopsidis* | Emoy2 | ERR012782-ERR012787, ERR015788, ERR015788 |
| *Hyaloperonospora* | *arabidopsidis* | Cala2 | SRR3254743 |
| *Hyaloperonospora* | *arabidopsidis* | Noks1 | SRR3254744 |
| *Peronospora* | *effusa* | R14 | SRR7107520 |
| *Phytophthora* | *agathidicida* | NZFS3770 | SRR2124796 |
| *Phytophthora* | *agathidicida* | NZF3772 | SRR2124795 |
| *Phytophthora* | *capsici* | Pc33E | SRR5436369 |
| *Phytophthora* | *capsici* | PC1E8 | SRR5436366 |
| *Phytophthora* | *infestans* | P3683 | SRR1817202 |
| *Phytophthora* | *infestans* | PIC99189 | ERR1990236 |
| *Phytophthora* | *nicotianae* | R0 | SRR2245478 |
| *Phytophthora* | *nicotianae* | R1 | SRR2198696 |
| *Phytophthora* | *parasitica* | P1569 | SRR653623 |
| *Phytophthora* | *parasitica* | P. Nic Tob race 0 | SRR653713 |
| *Phytophthora* | *parasitica* | P10297 | SRR714753 |
| *Phytophthora* | *parasitica* | P1976 | SRR653714 |
| *Phytophthora* | *ramorum* | Pr-106 | SRR6015292 |
| *Phytophthora* | *ramorum* | WSU115-0118 | SRR6015284 |
| *Phytophthora* | *ramorum* | WSU108-0021 | SRR6015282 |
| *Phytophthora* | *ramorum* | TKTK001E | SRR2141229 |
| *Phytophthora* | *ramorum* | TKTK001C | SRR2141227 |
| *Phytophthora* | *ramorum* | CC1048 | SRR4309669 |
| *Phytophthora* | *sojae* | P7076 | SRR1046800 |
| *Phytophthora* | *sojae* | ACR10 | SRR1046799 |
| *Pythium* | *arrhenomanes* | PAR_AB | SRR234858 |
| *Pythium* | *irregulare* | CBS 250.28 | SRR234877 |
| *Pythium* | *iwayami* | P174 | SRR235482 |
| *Pythium* | *ultimum var. sporangiiferum* | CBS219.65 | SRR235484 |
| *Saprolegnia* | *diclina* | VS20 | SRR576639 |
| *Saprolegnia* | *parasitica* | N12 | SRR515950 |
| *Saprolegnia* | *parasitica* | CBS223.65 | SRR058717 |
| *Sclerospora* | *graminicola* | Sg-4 | DRR097254 |
| *Sclerospora* | *graminicola* | UoM-SG-Pathotype1 | SRR3658180 |
